# Can machine learning automatically detect the aligned trunk in sitting directly from raw video using a depth camera?

**DOI:** 10.1101/421172

**Authors:** Maria Beatriz Sanchez, Ryan Cunningham, Penelope B Butler, Ian Loram

**Affiliations:** Manchester Metropolitan University

**Keywords:** Alignment, Objective measure, Machine learning, Deep convolutional neural network, Depth camera

## Abstract

**Background:** The Segmental Assessment of Trunk Control (SATCo) evaluates sitting control at seven separate trunk segments, making a judgement based on their position in space relative to a defined, aligned posture. SATCo is in regular clinical and research use and is a Recommended Instrument for Cerebral Palsy and Spinal Cord Injury-Paediatric by The National Institute of Neurological Disorders and Stroke (US). However, SATCo remains a subjective assessment.

**Research question:** This study tests the feasibility of providing an objective, automated identification of frames containing the aligned, reference trunk posture using deep convolutional neural network (DCNN) analysis of raw high definition and depth (HD+D) images.

**Methods:** A SATCo was conducted on sixteen healthy male adults and recorded using a Kinect V2. For each of seven segments tested, two different trials were collected (control and no-control) to simulate a range of alignment configurations. For all images, classification of alignment obtained from a trained and validated DCNN was compared to expert clinician’s labelling.

**Results:** Using leave-one-out testing, at the optimal operating threshold, the DCNN correctly classified individual images (alignment v misaligned) with average precision 92.7±16% (mean±SD).

**Significance:** These results show for the first time, automation of a key component of the SATCo test, namely identification of aligned trunk posture directly from raw images (HD+D). This demonstrates the potential of machine learning to provide a fully automated, objective SATCo test to enhance assessment of trunk control in children and adults for research and treatment of various conditions including neurodisability and stroke.

## 1. Introduction

Maintaining an upright posture of the head and trunk (trunk) is essential for humans to be able to perform everyday activities. Our ability to sit or to stand independently (i.e. without back or upper limb support) typically develops during our first year of life. Children with neurodevelopmental disorders, such as cerebral palsy or neuromuscular dystrophies, or adults with neurological conditions such as a stroke, frequently have compromised ability to sit independently, leading to functional limitations (Jensen and van Zandwijk, 2012; van Nes et al., 2008; Wee et al., 2015).

Evaluating trunk control in these children and adults is of paramount importance when planning therapeutic interventions that aim towards independent, unsupported functional sitting ability with a vertically aligned (neutral) head and trunk posture (Butler et al., 2010; Saeys et al., 2011; van Nes et al., 2008; Wee et al., 2015). Current physical therapy assessments are generally based on tests that evaluate trunk control status from the observation of functional abilities (Heyrman et al., 2011; Perlmutter et al., 2010; Pountney et al., 1999; Reid, 1997; Russell et al., 2002) or from balance tests in sitting (Verheyden et al., 2004). These assessments, although reliable, are subjective and most consider the head/trunk as a single unit, ignoring both its multi-segmental composition and any use of the hands for support to maintain a balanced posture (Heyrman et al., 2011; Pountney et al., 1999; Reid, 1997; Russell et al., 2002; Verheyden et al., 2004). The Segmental Assessment of Trunk Control (SATCo) is unique in addressing these issues, evaluating control based on 1) the position of individual trunk segments in space relative to a defined aligned posture and 2) the use of external support (Butler et al., 2010). SATCo is in regular clinical and research use and is a Recommended Instrument for Cerebral Palsy and Spinal Cord Injury-Paediatric by The National Institute of Neurological Disorders and Stroke (US); however, it currently is also a subjective assessment.

The generation of fully automated tools that are feasible for use in a clinical environment, will positively impact the planning and delivery of therapeutic interventions both for children and adults. Objective tools will provide a reliable and accurate evaluation of trunk control: this will help to quantify the changes due to therapy and the benefits of specific therapeutic interventions or combinations of therapies. Previous work has established that objective replication of the SATCo could be made (Sánchez et al., 2017a, b; Sánchez et al., 2018) but had the disadvantage of requiring a large amount of manual processing of video-based recordings.

In previous work, we have demonstrated a neural network methodology to automate tracking of individual trunk segments in seated children, comprising the seven cervico-thoraco-lumbar segments required for the Segmental Assessment of Trunk Control (SATCo) (Cunningham et al., 2018). While this work (Cunningham et al., 2018) demonstrates, for the first time, a feasible technical solution to automate tracking of individual trunk segments in a given sitting posture, and of changes away from that posture, it leaves unsolved the automated identification of the aligned trunk posture to act as a reference.

In this preliminary work we test a methodology to detect the reference aligned seated trunk posture for each participant to determine whether a fully powered study along these lines would be likely to lead to a successful system. Since SATCo requires estimation of the absolute posture of all segments relative to the reference aligned posture, it is important to minimise err
or in the estimated reference posture. The method tested here is required to provide, for each participant, a robust estimate of one or more frames containing the aligned trunk posture from which a segmental reference posture could be calculated.

Specifically, we test whether a trained neural network can detect automatically the aligned trunk posture in sitting directly from raw video from a Kinect V2 camera during SATCo testing of healthy adult males. State of the art machine learning methods were applied to predict the clinical labels (frames classified as aligned or not aligned) (Radford et al., 2015; Zeiler et al., 2010). This enabled examination of whether automated analysis of raw high definition and depth (HD+D) images is suitable for the identification of aligned frames and selection of frames that best represent the aligned trunk posture in sitting by comparing the expert’s labelling to the machine learning classification.

## 2. Methods

Ethical approval was obtained from the Manchester Metropolitan University Ethics Committee. The participants were 16 healthy adult males (mean±SD age 31.39 ±5.21 years, mean±SD height 1.78m ±0.07, and weight 77.7kg ±11.1). All participants provided written informed consent to the work. Participants sat on a bench free of back or arm support; the height of the bench ensured that participants’ feet were flat on the floor and the knees and hips were flexed at 90o. A Kinect V2 camera was set at a distance of 1.60m and height of 0.90m on the left side of the participants and both grayscale HD and depth images were recorded at 15Hz (asynchronous recording software was written by the authors in c++). The SATCo was conducted; a tester provided manual support to the participant’s trunk to test seven discrete trunk segmental levels following the published guidelines (Butler et al., 2010). Two different trials were collected, control and no-control, to simulate a range of alignment configurations. For the control trials, participants were asked to remain still for 5s in upright sitting with the arms and hands free in the air; for the no-control trials a verbal cue was given for participants to simulate lack of trunk control by making movements of the unsupported segments of the trunk (segments above the tester’s hand support) away from the aligned position (e.g. falling forwards). All segmental levels were included within the analysis.

Figure 1 shows the neural network architecture. The analysis steps were as follows:

**Figure 1.**
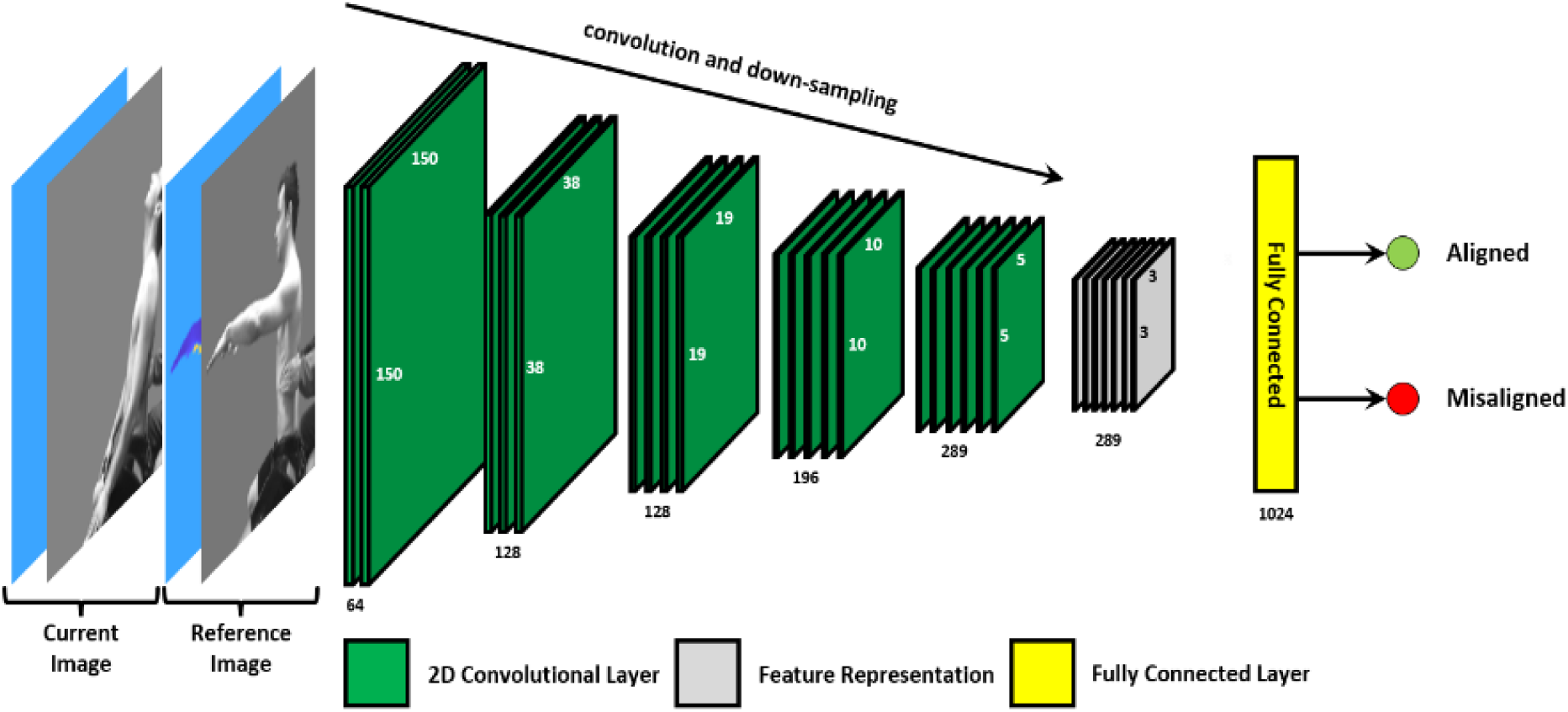
Network architecture. Showing the deep convolutional neural network (DCNN) architecture consisting of 2 input channels (depth and grey-converted RGB image), 5 convolutional and down-sampling (max-pooling) layers, followed by a fully connected layer, and a logistic regression output layer.

i. Training labels (classification of each image) for the neural network were provided by a clinician (MBS) with four years of experience doing this analysis. MBS identified the frames when all unsupported segments in the trunk were aligned, and the frames when one or more unsupported segments were misaligned.
ii. Using raw depth information from the Kinect, the background was automatically subtracted from the image to reveal the participant and tester.
iii. Eight identical multilayer deep convolutional neural networks (DCNN) were trained to provide held-out test results for all 16 participants. Each network used approximately 22,000 images from 14 participants and each tested on approximately 3,000 images from two participants not used for training.
iv. The neural network was trained to predict the clinical labels, from an individual image at time t.
v. After training, for all 16 participants, the neural network output varying 0 to 1 (aligned) was compared to the clinical labels using precision – recall analysis. Using leave-one-out cross validation (LooCV), an optimal neural network prediction threshold was selected to maximise the precision (ratio of true aligned to predicted aligned classification).
vi. We investigated whether precision was improved by excluding frames at the transition between predicted aligned and predicted non-aligned states.

N.B. Precision is the fraction of frames predicted aligned, which are correct. Recall is the fraction of aligned frames which are predicted aligned (true positive rate). “Average Precision” (AP) reports the mean precision over the entire precision-recall curve, with frames predicted as aligned sorted by descending neural network prediction score. “Precision” reports precision at maximum recall.

## 3. Results

The neural network appears to classify frames correctly for trunk alignment (Figure 2). Comparison with labels showed the prediction threshold made little difference to the precision of the results (Figure 3). By removing frames at the transition between aligned and non-aligned states, AP (averaged for all participants) improved from 84.1% with no transitional frames excluded to a maximum of 94.7% at prediction threshold 0.99 and transition time 0.8s. Average precision was maintained robustly over 89.5% for a broad range of parameters, namely all thresholds less than 0.994 and transition times of 0.5 to 1.7s. For each participant, using all other participants to select the optimal threshold and transition time, the neural network classified frames (alignment v. misalignment) with AP of 92.7 ± 16% (mean±SD) (Table 1, Figure 3).These results include low accuracy for participant 9, (AP 36%), and high accuracy generally indicated by the median for all participants of AP 100% and a second lowest AP of 80.5% (Table 1, Figure 3).

**Table 1.** Summary of results. This table presents comprehensive results for each participant, calculated over all trials, and a summary of results, calculated over all participants and trials. For each participant, the number of positively classified (*aligned*) and negatively classified (*misaligned*) images are given for the clinical expert and for the automated system, followed by measures of agreement between the clinical expert and the neural network (ANN), in the form of accuracy, precision, recall, F1 score (harmonic mean of precision and recall), and average precision. Values are calculated using LooCV. For each participant, the threshold and transition time was selected to give the maximum averaged precision on all other participants.

| Participant # | # Positive classes (labels) | # Negative classes (labels) | True Positive rate (TPr) | False Positive rate (FPr) | True Negative rate (TNr) | False Negative rate (FNr) | Precision (%) | $F_1$ (%) | Average Precision (AP) |
| --- | --- | --- | --- | --- | --- | --- | --- | --- | --- |
| 1 | 693 | 257 | 93.5 | 17.9 | 82.1 | 6.5 | 93.4 | 93.4 | 93.5 |
| 2 | 894 | 73 | 0.6 | 4.1 | 95.9 | 99.4 | 62.5 | 1.1 | 94.3 |
| 3 | 441 | 186 | 32.7 | 4.8 | 95.2 | 67.3 | 94.1 | 48.5 | 93.1 |
| 4 | 671 | 22 | 100.0 | 0.0 | 100.0 | 0.0 | 100.0 | 100.0 | 100.0 |
| 5 | 767 | 68 | 15.3 | 0.0 | 100.0 | 84.7 | 100.0 | 26.5 | 100.0 |
| 6 | 591 | 240 | 5.9 | 0.0 | 100.0 | 94.1 | 100.0 | 11.2 | 100.0 |
| 7 | 734 | 309 | 4.6 | 0.0 | 100.0 | 95.4 | 100.0 | 8.9 | 100.0 |
| 8 | 612 | 177 | 16.2 | 0.0 | 100.0 | 83.8 | 100.0 | 27.8 | 100.0 |
| 9 | 196 | 406 | 98.0 | 51.0 | 49.0 | 2.0 | 48.1 | 64.5 | 36.3 |
| 10 | 493 | 80 | 47.1 | 0.0 | 100.0 | 52.9 | 100.0 | 64.0 | 100.0 |
| 11 | 557 | 35 | 100.0 | 0.0 | 100.0 | 0.0 | 100.0 | 100.0 | 100.0 |
| 12 | 688 | 179 | 81.3 | 41.9 | 58.1 | 18.8 | 88.2 | 84.6 | 98.8 |
| 13 | 729 | 91 | 14.8 | 0.0 | 100.0 | 85.2 | 100.0 | 25.8 | 100.0 |
| 14 | 383 | 105 | 38.6 | 47.6 | 52.4 | 61.4 | 74.7 | 50.9 | 80.5 |
| 15 | 782 | 199 | 3.3 | 5.0 | 95.0 | 96.7 | 72.2 | 6.4 | 100.0 |
| 16 | 332 | 126 | 90.7 | 22.2 | 77.8 | 9.3 | 91.5 | 91.1 | 86.2 |
| Mean | 598 | 160 | 46.4 | 12.2 | 87.8 | 53.6 | 89.1 | 50.3 | 92.7 |
| SD | 187 | 106 | 40.2 | 18.5 | 18.5 | 40.2 | 16.1 | 35.9 | 16.1 |
| Median | 401 | 36 | 95.5 | 57.9 | 42.1 | 4.5 | 96.8 | 92.3 | 99.2 |

**Figure 2.**
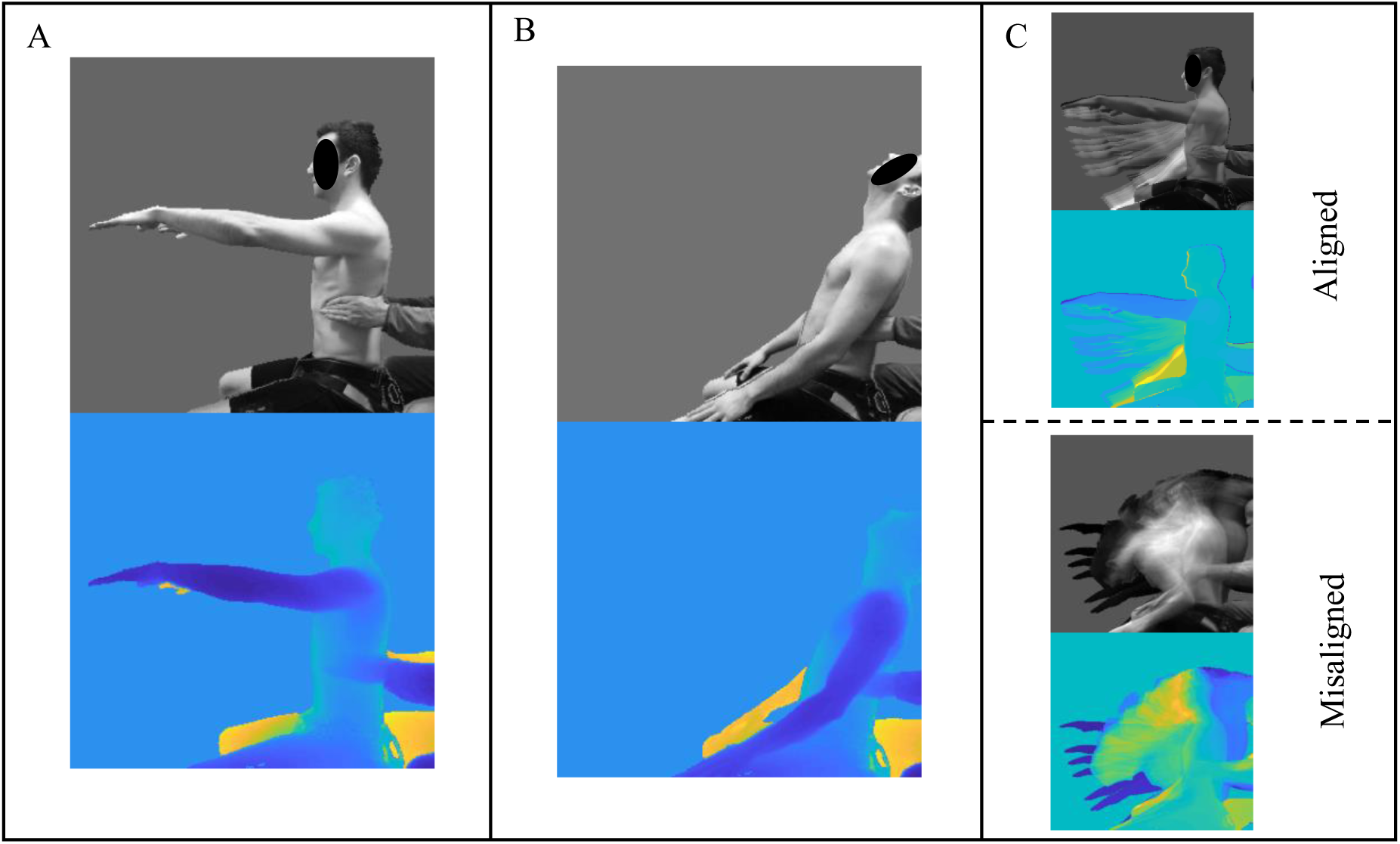
Representative trial example of Static control when testing at the Lower-Thoracic segmental level. In all grayscale and depth images, the background was subtracted using the information in the depth image. Showing **A)** an example of ‘aligned’ trunk posture (classified by the expert) presented in the form of grayscale (top) and depth image (bottom); **B)** an example of ‘misaligned’ trunk posture (classified by the expert) presented in the form of grayscale and depth image; and **C)** distribution of grayscale and depth images, correctly classified by the neural network as ‘aligned’ (top) or ‘misaligned’ (bottom).

**Figure 3.**
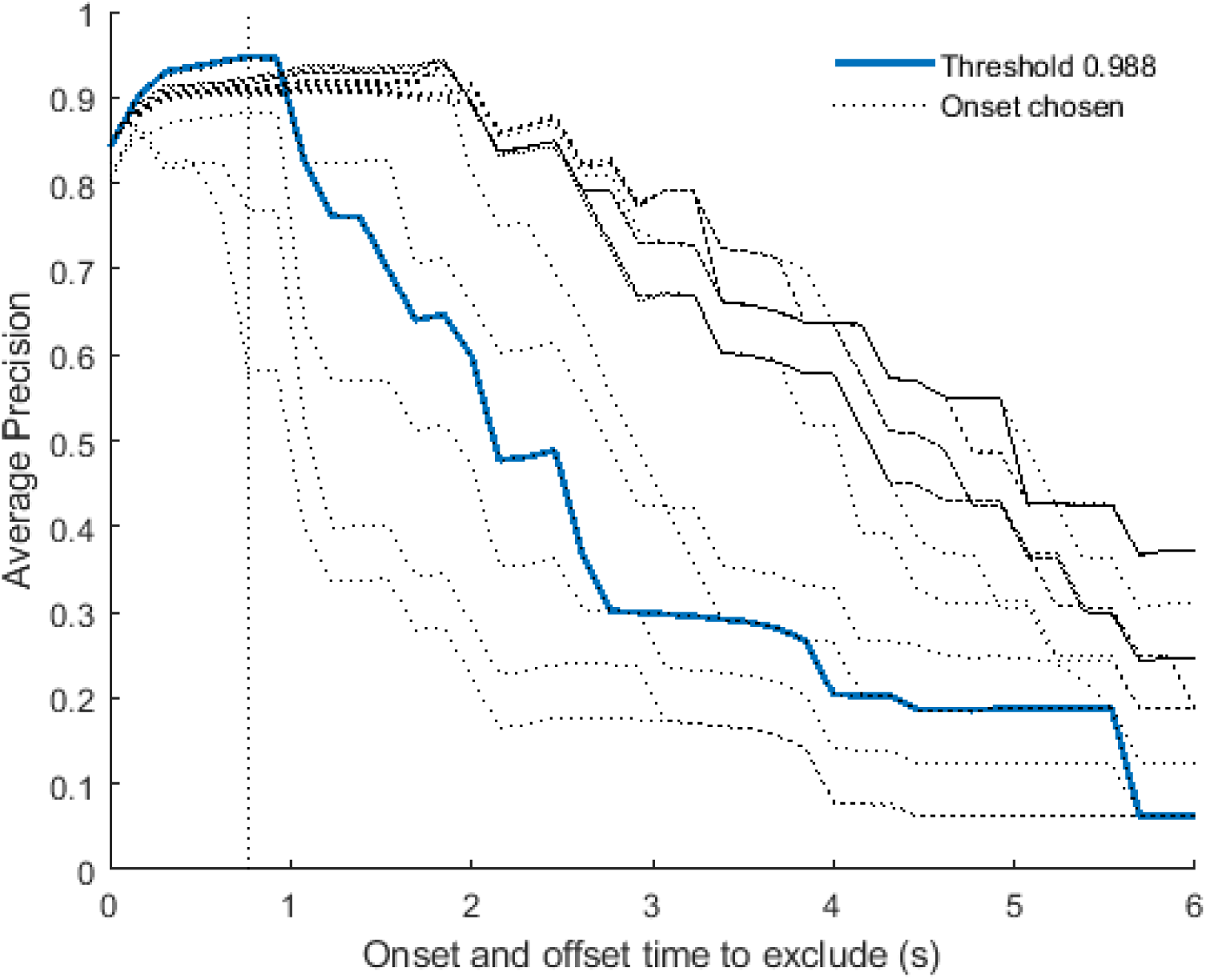
Variation of Precision with threshold and exclusion of transitional frames. Panel shows “Average Precision” (AP) v duration of frames excluded from the start and from the end of any sustained aligned or sustained non-aligned state. AP is averaged across all 16 participants. Blue solid line is threshold 0.988. Faint dashed lines show other thresholds (1.00, 0.996, 0.994, 0.992, … 0.6 running lower left to upper right). Vertical dashed line (0.77s) is median transition time chosen for results in Table 1.

## 4. Discussion

This study tested a neural network methodology to provide automated identification of frames containing the aligned trunk posture for a sitting person. The main requirement for use within an automated SATCo assessment is that frames identified as aligned, are indeed aligned. Average Precision (AP) is the measure of performance reporting this requirement. Frames containing the aligned trunk posture will be used to calculate a participant reference posture that will be used for all trials within a SATCo test. A full SATCo test of up to seven segments, including Static, Active and Reactive could include 20 trials (Butler et al., 2010). With a median AP of 100%, and AP below 80% for only one participant (Table 1), these results demonstrate an encouraging level of success suggesting that a fully powered study along these lines on clinical populations would be likely to lead to a successful system.

This method used a single Kinect camera and an analysis that is markerless. The results (Figure 2) depict the quality of data that can be collected with this method, the ability for automatic removal of background to reveal the participant showing that there is very little noise in both images, and to differentiate the left and right limbs. These features aid correct classification. Clinical labels were required only for training; thereafter the neural network provided fully automatic analysis of alignment without any further user interaction.

Participants simulated lack of trunk control by moving away from the aligned posture; movement displacement occurred in all three planes of motion. Previous studies used a 2D video-based semi-automated method to track markers in the sagittal plane (Sánchez et al., 2017a; Sánchez et al., 2018); participant’s movement in planes other than the sagittal generated movement artefacts reducing the accuracy of the method. The method reported here enabled the accurate classification of trunk alignment from raw HD+D images even in situations where motion occurred orthogonal to the sagittal plane. This methodology to identify a reference posture is intended to be used in conjunction with a parallel process providing automated tracking of specific landmarks of the head, trunk and arms in children with cerebral palsy to identify which segments are misaligned (Cunningham et al., 2018) and where support is gained through the upper limbs. For all frames identified as containing an aligned trunk posture, the point features would be averaged to provide a segmental estimate (mean±SD) of the reference trunk posture (Sánchez et al., 2017a; Sánchez et al., 2018). The absolute angle of individual trunk segments would be calculated relative to the referenced aligned posture.

Classification of trunk alignment in sitting is one of the components required to assess control status in the Segmental Assessment of Trunk Control (SATCo). This was a preliminary study of a non-clinical population mimicking control/no-control. Nevertheless, the results demonstrate proof of principle that this deep convolutional neural network (DCNN) can automate the objective classification of trunk alignment in sitting from raw HD+D images. This proof of concept justifies development of the neural network analysis to the accuracy required for clinical use. Such a tool would enhance the understanding of the typical development of head and trunk control and its alterations in the presence of neuromotor disabilities (Jensen and van Zandwijk, 2012) and of the role of trunk control in recovery from stroke (van Nes et al., 2008; Wee et al., 2015).

## Acknowledgements

With appreciation, we thank the participants who gave their time for this study and Brian Bate who assisted with the implementation of SATCo procedures during data collection..

